# Recombination suppression in heterozygotes for a pericentric inversion induces interchromosomal effect on crossovers in *Arabidopsis*

**DOI:** 10.1101/376376

**Authors:** Pasquale Termolino, Matthieu Falque, Gaetana Cremona, Rosa Paparo, Antoine Ederveen, Olivier C. Martin, Federica M. Consiglio, Clara Conicella

## Abstract

During meiosis, recombination ensures the allele exchange through crossovers (COs) between the homologous chromosomes and, additionally, their proper segregation. CO events are under a strict control but molecular mechanisms underlying CO regulation are still elusive. Some advances in this field were made by structural chromosomal rearrangements that are known at heterozygous state to impair COs in various organisms. In this paper, we have investigated the effect that a large pericentric inversion involving chromosome 3 of *Arabidopsis thaliana* has on male and female recombination. The inversion associated to a T-DNA dependent mutation likely resulted from a side effect of the T-DNA integration. Reciprocal backcross populations, each consisting of over 400 individuals, obtained from the T-DNA mutant and the wild type, both crossed with *Landsberg*, have been analyzed at genome-wide level by 143 SNPs. We found a strong suppression of COs in the rearranged region in both male and female meiosis. As expected, we did not detect single COs in the inverted region consistently with the post-meiotic selection operating against unbalanced gametes. Cytological analysis of chiasmata in F1 plants confirmed that COs are effectively dropping in chromosome 3 pair. Indeed, CO failure within the inversion is not altogether counterbalanced by CO increase in the regions outside the inversion on chromosome 3. Strikingly, this CO suppression induces a significant increase of COs in chromosome pairs 1, 2 and 5 in male meiosis. We conclude that these chromosomes acquire additional COs thereby compensating the recombination suppression occurring in chromosome 3, similarly to what has been described as interchromosomal (IC) effect in other organisms. In female meiosis, IC effect is not evident. This may be related to the fact that CO number in female is close to the minimum value imposed by the obligatory CO rule.

**Author Summary:** It is well known that chromosome structure changes in heterozygous condition influence the pattern of meiotic recombination at broad scale. In natural populations, inversions are recognized as the most effective force to reduce COs. In this way, adaptive allele combinations which otherwise would be broken by recombination are maintained. In the present work, we studied the effect on recombination of a large pericentric inversion involving Arabidopsis chromosome 3. The analysis on heterozygous populations provided evidence of strong recombination suppression in chromosome 3. However, the most striking aspect of this study is the finding that the failure of chromosome 3 to recombine is coupled to increased CO frequencies on the other chromosome pairs in male meiosis. These CO compensatory increases are strictly an interchromosomal (IC) effect as was first described in *Drosophila*. As far as we know, it is the first time IC effect has been reported in plants. Unfortunately, the molecular mechanisms underlying IC effect in the other organisms are still elusive. To understand how a CO change on just one chromosome triggers the global response of the meiocyte to obtain the adequate CO number/cell remains a fascinating question in sexually reproducing species.

## Introduction

Meiotic recombination is fundamental in creating genetic diversity through the shuffling of alleles over generations. Recombination is initiated by the formation of DNA double-strand breaks (DSBs) that are repaired, partly, as crossovers (COs), i.e., through reciprocal exchanges between the homologous chromosomes. Since more DSBs are generally formed than COs, alternative pathways of DSB repair occur as non-crossover (NCO), i.e. nonreciprocal exchange, and inter-sister repair (IS) [1]. CO rate is suggested to be a trait under selection for lower as well as upper limits [2]. Effectively, CO formation is bounded by numerous constraints. Indeed, to ensure proper chromosome segregation at Meiosis I, at least one CO per homologous chromosome pair (obligatory CO) has to occur while the formation of excess COs is almost always limited, possibly by mechanical reasons (such as entanglement) or evolutionary pressures (putative mutagenicity of COs). The complex CO control system includes CO homeostasis that maintains nearly constant CO number per meiosis despite variation in DSB numbers [3] and CO interference that leads COs to be along the chromosome farther apart than expected by chance [4]. The molecular mechanisms underlying CO homeostasis and interference are still elusive. So far, CO homeostasis has been documented in yeast [3] and mouse [5]. In the latter, the authors suggested that homeostatic control operates at the different steps of meiotic recombination in a progressive manner, i.e., from the formation of early recombination intermediates till CO formation after homologs have fully synapsed during pachytene. In yeast, a negative feedback loop mechanism is thought to impose the formation of additional DSBs until DSB number is sufficient to obtain an adequate CO rate [6]. As reported recently, CO homeostasis does not seem to be prominent in maize [7] suggesting that CO homeostatic control is not strong in plants. Conversely, the majority of COs is subject to strong interference in plants. For example, in *Arabidopsis* about 85% of total COs exhibit interference [8].

Mutations in components of structures or pathways limiting COs allow to overcome the upper limit of CO number typical of a given organism. For example, partial depletion of proteins of the synaptonemal complex central region increases COs in *C. elegans* [9]. In *Arabidopsis*, the disruption of pathways that drive DSB repair towards NCOs induces a boost in CO rate by almost eight fold, apparently without any negative consequences [10]. On the other hand, the lower limit of CO number, referred to as ‘CO assurance’, has to be respected since at least one CO per bivalent is necessary to avoid abnormal disjunction at anaphase I and unbalanced gametes. In natural populations, mechanisms decreasing COs (but not below the lower limit) are tolerated probably because they maintain adaptive allele combinations which otherwise would be broken by recombination [11]. One of these mechanisms operates through inversions, which are strong suppressors of recombination across the rearranged chromosomal regions [12]. Reduction/suppression of COs in inverted regions is reported in various organisms including yeast [13], *C. elegans* [14], mammals [15, 16] and plants [17–19]. Inversions have been studied extensively in multiple *Drosophila* species [20]. In particular, heterozygous inversions suppressing COs there also enhance the frequency of COs on the remaining chromosome pairs in a phenomenon referred to as the *‘interchromosomal effect’* [21], a nomenclature that will be used hereafter.

In this paper, we investigated the recombination rate in *Arabidopsis* heterozygous populations of the mutant *meiotic control of crossover 1 (Atmcc1)*. Male meiosis of *Atmcc1*, that was isolated from an enhancer activation tagging population [22], was previously characterized [23]. In the present work, we ascertained that a pericentric inversion was associated to the *Atmcc1* mutation, probably as side effect of T-DNA integration. Such chromosomal rearrangements are common at T-DNA integration sites [24]. By examining the effect of the inversion on CO rate in both male and female meiosis, we evidenced a CO decrease in the chromosome carrying the inversion and more interestingly the associated interchromosomal effect, similar to what has been described in *Drosophila* [21].

## Results

### *Atmcc1* mutation is associated with a structural rearrangement in chromosome 3

In this work, we investigated the recombination in male and female meiosis by analyzing *Atmcc1* heterozygous populations. We generated F1 progenies by crossing *Landsberg erecta* (Ler) with homozygous *Atmcc1* (Ler x *Atmcc1*) as well as Ler x C24 as control. Subsequently, we crossed these F1 plants, as male and female, with Ler thereby obtaining four large reciprocal backcross (BC1) populations (Mmut, Mctr, Fmut, Fctr) (Fig 1). The number of COs, revealed by means of 143 SNPs analyzed in the four BC1 populations, were 1871 in Mmut (N=418 plants) and 1259 in Fmut (N=417 plants), while 1834 and 1279 COs were found in Mctr (N=414 plants) and Fctr (N=410 plants), respectively. The total CO number is similar between the respective mut and ctr populations (Mmut *vs* Mctr; Fmut *vs* Fctr; p=0.5, Fisher’s test). Accordingly, the estimated average number of COs per cell is 8.95 *vs* 8.86 (Mmut *vs* Mctr) for male meiosis and 6.04 *vs* 6.24 (Fmut *vs* Fctr) for female meiosis (Table 1). On the other hand, CO number is significantly different between male and female meiosis, in agreement with the known heterochiasmy (p=0.0037, χ^2^ test). The recombination rate calculated as the ratio between the genetic map length in centiMorgan and the physical length in megabase pair was estimated by Marey maps for each chromosome (Fig 2, S1 Fig). The marker positions refer to Ler as reported in Weigel database [25, 26]. In control populations, CO number and distribution in male (Table 1, Fig 2) and female meiosis (Table 1, S1 Fig) are substantially in agreement with the results published by Giraut and colleagues [27] even though genotypes and markers used in our study are only in part the same. We confirmed that male recombination occurred predominantly near both telomeres whereas pericentromeric regions were almost devoid of COs. As expected in female meiosis, the regions close to telomeres showed a lower recombination with respect to male meiosis.

**Fig 1.**
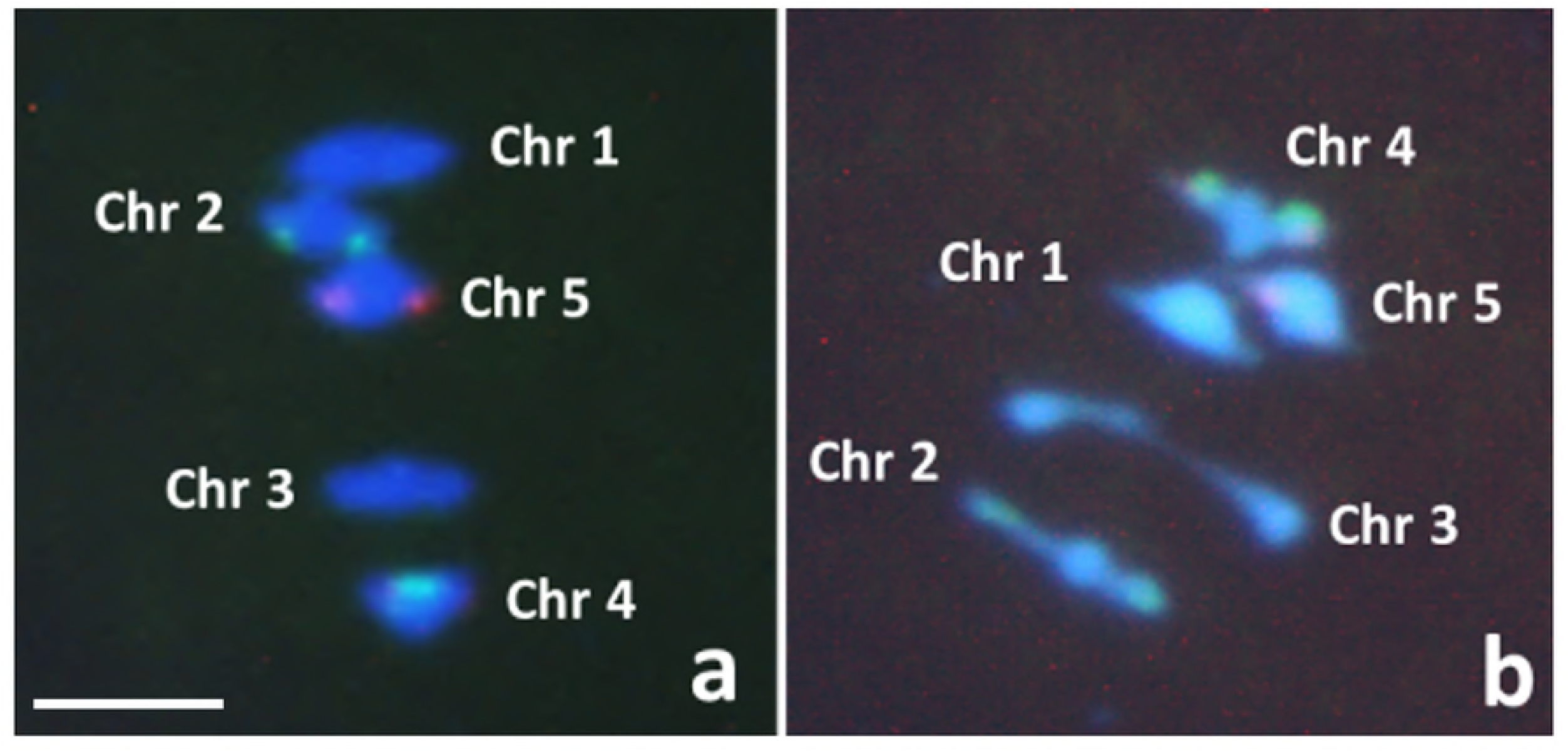
Schematic representation of the crosses performed to obtain the four reciprocal backcross (BC1) populations from *Atmcc1* mutant and wild type in Arabidopsis. Fctr, Mctr = BC1s from F1 wild type plants used as female and male parent, respectively. Fmut, Mmut = BC1s from F1 mutant plants used as female and male parent, respectively. Ler = *Landsberg erecta* (recurrent parent), C24 = background of *Atmcc1*.

**TABLE 1.** Number of crossing overs (COs) per chromosome pair per cell in backcross populations from control (Mctr, Fctr) and mutant (Mmut, Fmut).

|  | Mctr | Mmut | Fctr | Fmut |
| --- | --- | --- | --- | --- |
| CHR1 | 2.30 | 2.63 | 1.47 | 1.58 |
| CHR2 | 1.29 | 1.66 | 1.13 | 1.14 |
| CHR3 | 1.75 | 0.55 | 1.07 | 0.63 |
| CHR4 | 1.51 | 1.51 | 1.25 | 1.28 |
| CHR5 | 2.01 | 2.60 | 1.32 | 1.41 |
| ALL | 8.86 | 8.95 | 6.24 | 6.04 |

ALL: all five chromosomes pooled.

In both mutant populations, absence of recombination was strikingly evident in chromosome 3 (Fig 2, S1 Fig, S2 Fig, Table 1). We hypothesized a possible structural rearrangement to be the causative factor of the recombination suppression. Evidence for the presence of such a structural variation in the mutant comes from two sources. First, when constructing the genetic map *ab initio*, chromosome 3 led to two possible orders with nearly identical likelihoods, these orders differing by an inversion of the block of markers CH3-3 to CH3-23, interval that contains the centromere. Second, based on the heat-map showing pairwise two-point linkage of this chromosome, we see that there is a linkage much greater than expected in this chromosome particularly between the markers CH3-3 and CH3-23 (Fig 3). These facts suggest that CHR3-3 and CHR3-23 might be the closest markers to the boundaries of the pericentric inversion.

**Fig 2.**
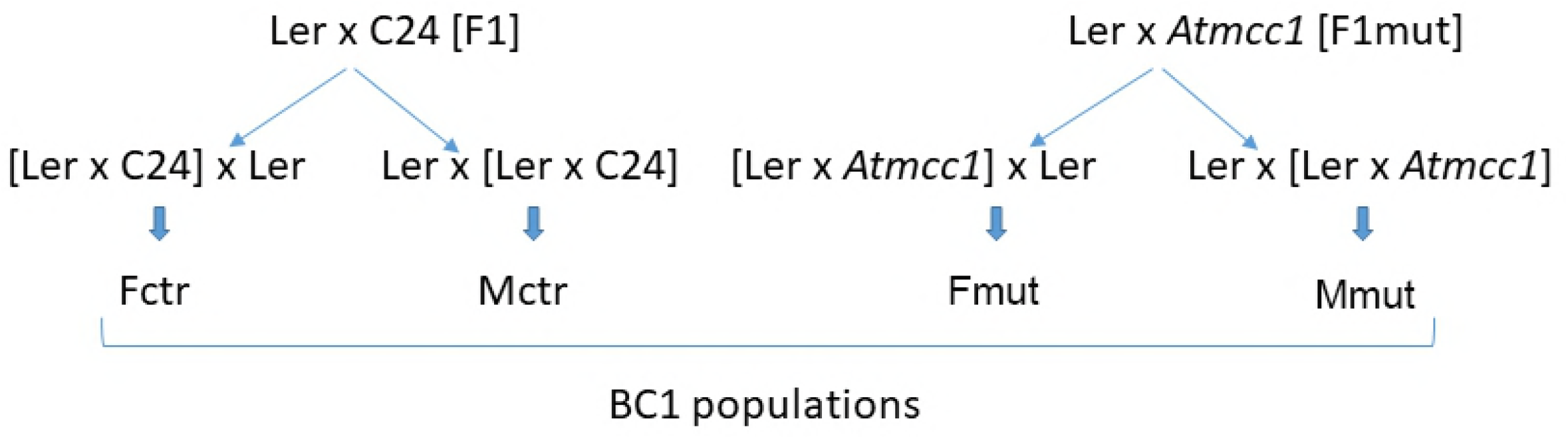
Relationship between physical and genetic positions, and corresponding recombination rate per chromosome. X-axis: physical position (Mbp) of the SNPs. Left Y-axis and blue lines: genetic position (cM) of SNPs on Mctr and Mmut linkage maps. Right Y-axis and red lines: recombination rate (cM/Mbp) given by the derivative of the smoothed Marey map curve.

**Fig 3.**
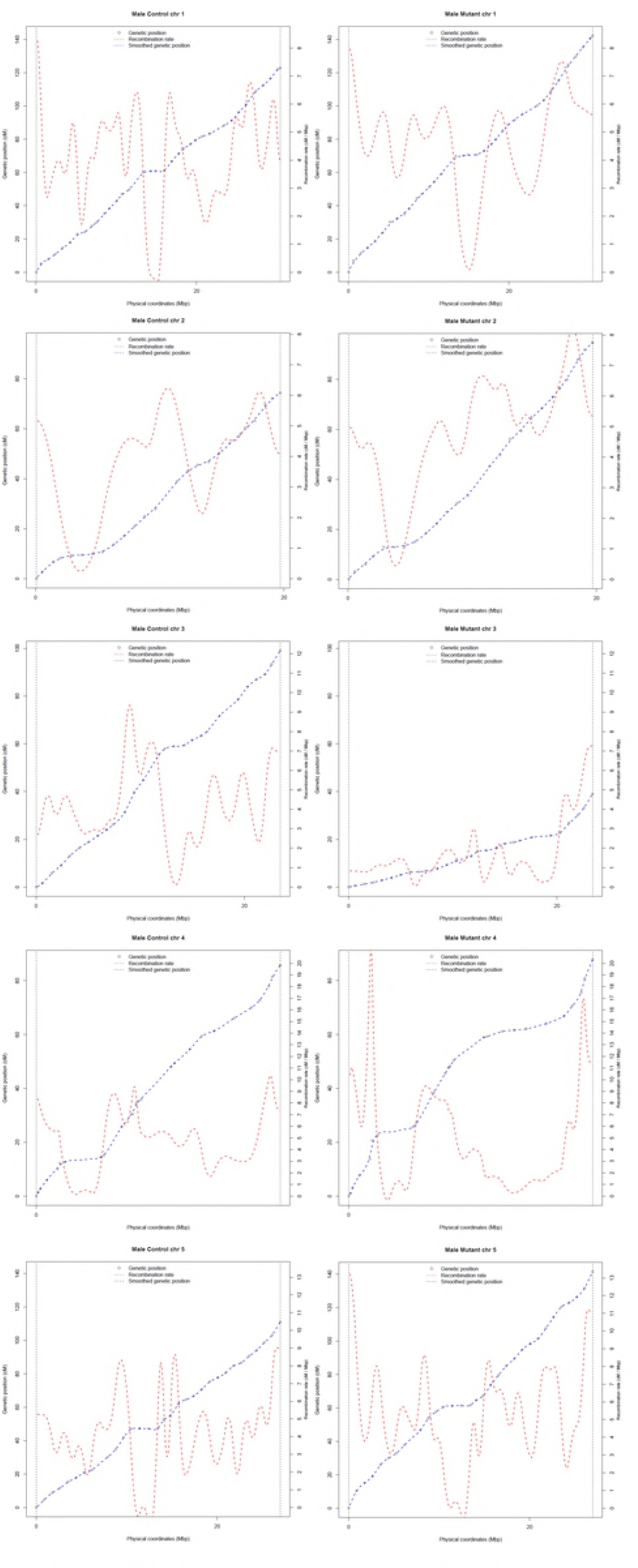
Genome-wide heat maps for pairwise genetic distances. Left: control population showing the standard structure of distances increasing as one goes away from the diagonal and no short distances between markers on different chromosomes. Right: mutant population showing the standard structure for chromosomes 1, 2 and 5 and abnormal behavior both within chromosome 3 and between chromosomes 3 and 4.

### Rearranged chromosome 3 leads to post-meiotic selection

As a proxy of the anticipated inverted region, hereafter we will assume that it is identical to the segment between markers CH3-3 and CH3-23 in the control. When examining the number of COs in this region, we find that the control has many cases of a single CO whereas the mutant almost never has a single CO in spite of having nearly as many double COs as the control (Fig 4). *A posteriori*, this is expected if an inversion is present, because an odd number of COs in that region produces two chromatids, each missing large genomic regions (Fig 5). These chromatids are eliminated by post-meiotic selection. Furthermore, an even number of COs within the inversion is expected to give viable products (Fig 5) and so cases of double COs should not be much affected. To sum up, a loss of genomic regions containing non-dispensable genes is expected to occur in the case of a single or odd number of COs within the inversion, thereby causing unviable post-meiotic products whereas in the cases of double or even number COs within the inversion, balanced and viable products are expected (Fig 5). COs forming exclusively in the regions outside the inversion are expected to have no effect on the viability of post-meiotic products (Fig 5). By keeping this in mind, we generated a working model of post-meiotic selection in *Atmcc1*. First, it is interesting to note that we found only products where the COs were in *just one* of the three regions: (1) markers CH3-1/CH3-2, (2) markers CH3-3/CH3-23, (3) markers CH3-24/CH3-27. For instance, individuals carrying simultaneously COs on both terminal regions outside the inversion were not observed. We have thus implemented this constraint into our model in addition to the constraint on having an even number of COs in the inversion. We take as unknown the pairing probabilities of the three regions. For simplicity, no interference was considered. The three parameters of our model were then adjusted to get agreement with the frequencies of the plants effectively observed in the different classes after post-meiotic selection. The data for male meiosis led to pairing probabilities of 0.08, 0.17, and 0.75 in regions 1, 2 and 3, respectively (data used to construct Fig 6). Similar results were found for female meiosis.

**Fig 4.**
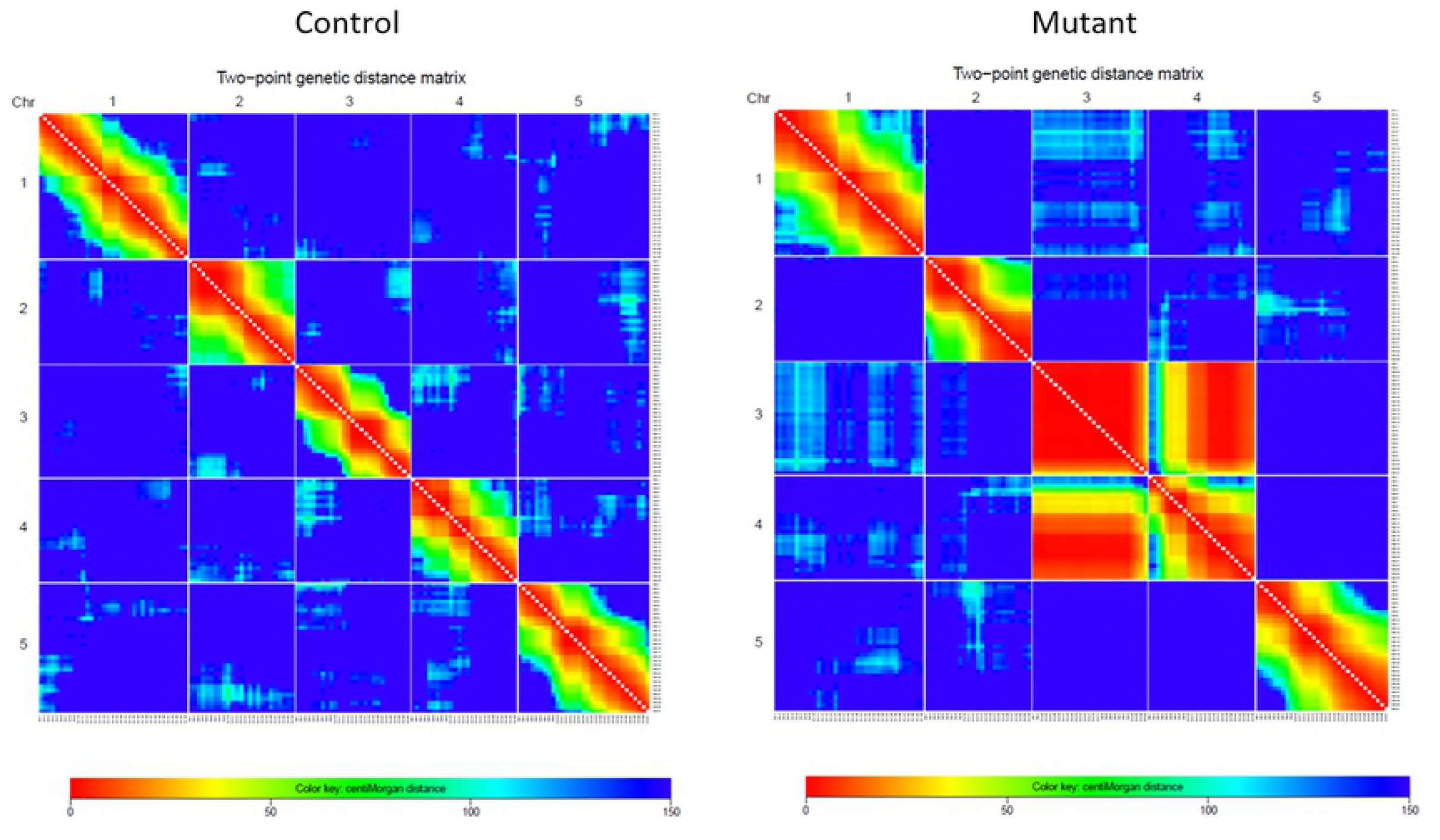
Number of single and double COs occurring in chromosome 3 region spanning CH3-3 and CH3-23 markers in the recombinant populations.

**Fig 5.**
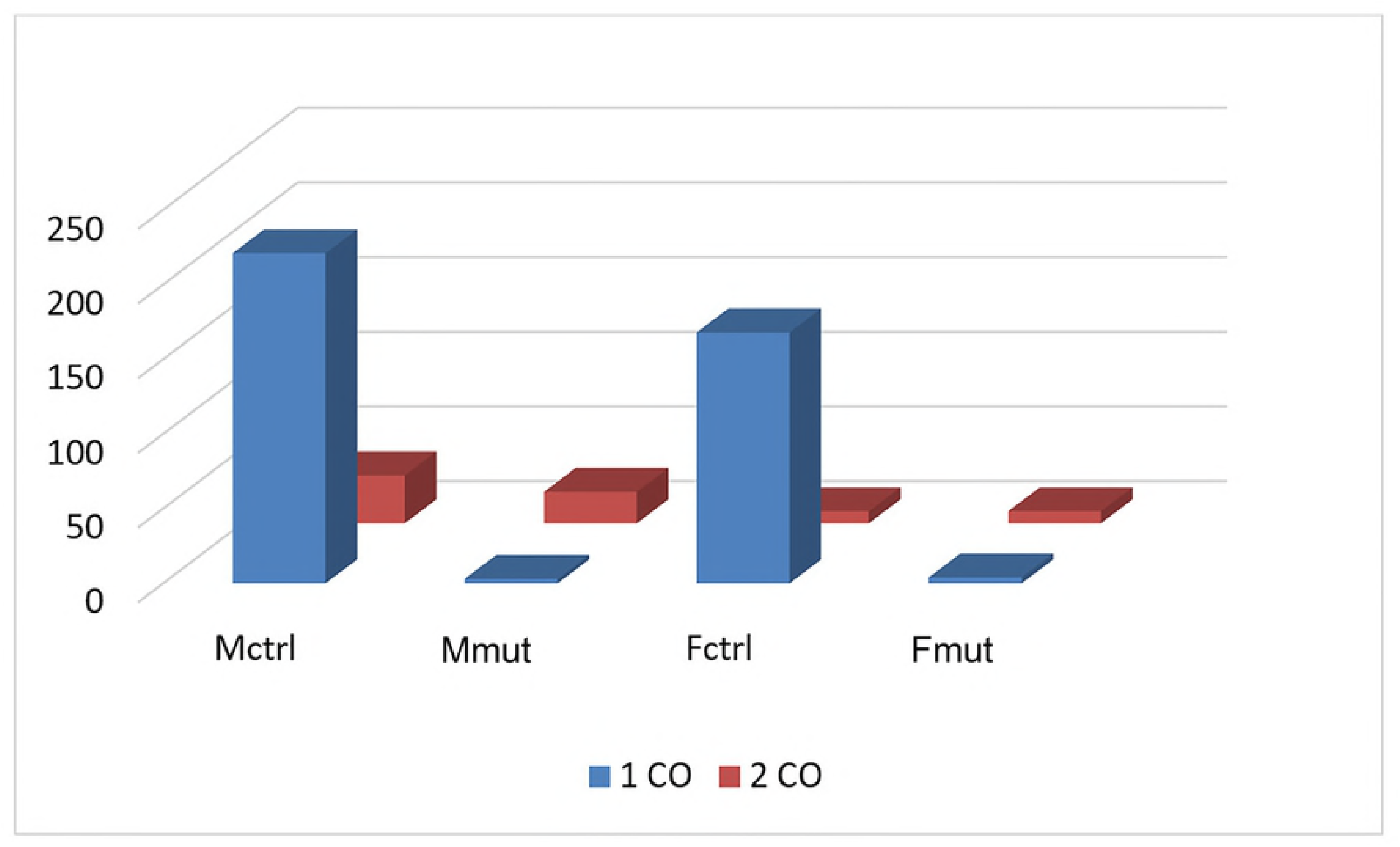
Schematic representation of pairing between homologous chromosome 3 differing for a pericentric inversion. Upper: Pairing of only one region outside the inversion without loop formation. Bottom: Pairing of the chromosomal region with inversion in the anti-parallel orientation (homo-synapsis) associated with the pairing failure of the outside regions. Even numbers of COs within inverted region produce viable gametes whereas odd numbers give unviable gametes. COs occurring in outside regions do not influence the gamete viability.

**Fig 6.**
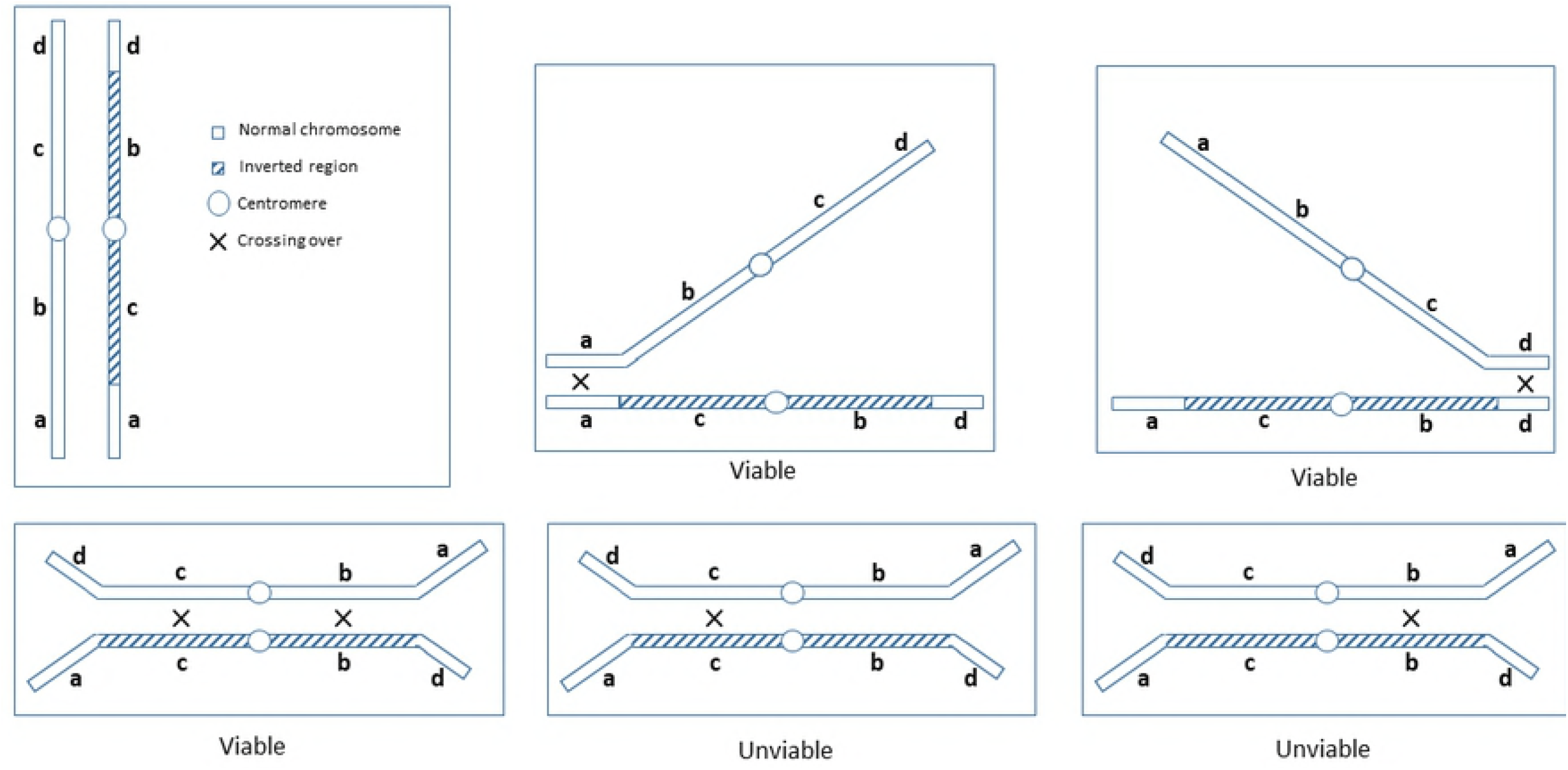
Heat maps for the two-point linkage LOD scores predicted by our model of the consequences of chromosome 3 inversion. Left: case simulating crossover formation and post-meiotic selection in male meiosis. Right: the same for female meiosis. Note the characteristic block structure of these heat maps in which the boundaries of the inverted region are being tightly linked (high LOD score).

### Rearranged chromosome 3 has suppressed recombination

Because of post-meiotic selection, genetic data cannot provide an estimation of CO number for chromosome 3 in *Atmcc1*. For this reason, we evaluated the number of chiasmata at metaphase I of male meiosis in F1 Ler x *Atmcc1* in comparison to F1 Ler x C24 (as control) according to the cytological method described by Sanchez-Moran and colleagues [28]. Chromosome 3 evidenced a significant decrease in the number of chiasmata per cell in Ler x *Atmcc1* vs control (1.11 *vs* 1.69, Fisher’s test <0.001) (Fig 7). Analysis of bivalent configurations revealed an increase of rod bivalents (where the chiasma is restricted to a single arm) in Ler x *Atmcc1* (88%; n=60 cells) when compared to Ler x C24 (30%; n=36 cells) (χ^2^ with Yates correction=2.268; p=0.01).

**Fig 7.**
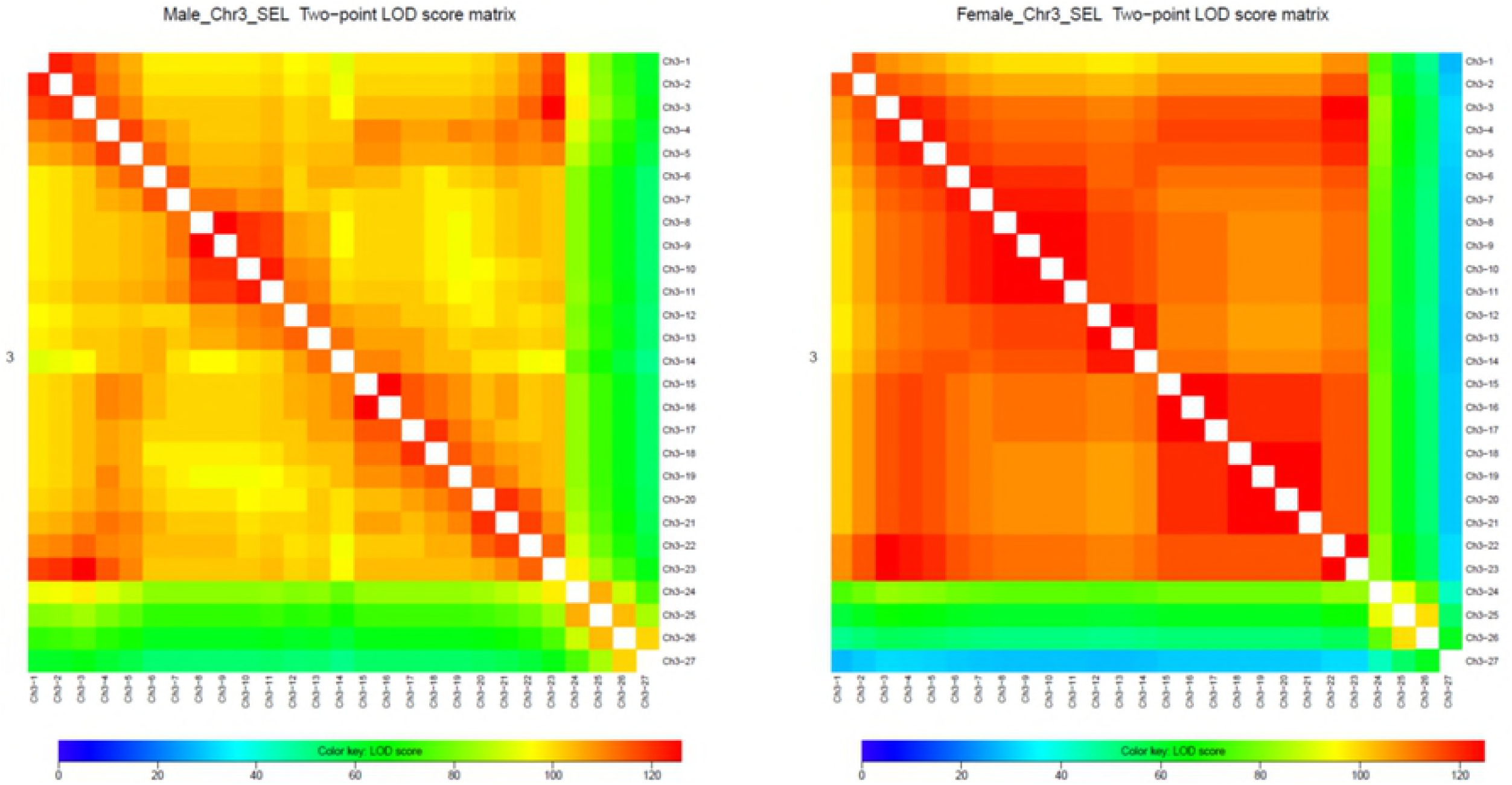
FISH from F1 cross Ler x C24 (a) and Ler x *Atmcc1* (b) at metaphase I in male meiosis. FISH is performed with 5S rDNA (red) and 45S rDNA (green) probes. DAPI stains DNA (blue). Scale bar: 5 μm.

### *Atmcc1* mutation is also associated with a structural rearrangement between chromosomes 3 and 4

Although the *ab initio* construction of individual genetic maps gave the same order as the physical one for chromosomes 1, 2, 4, and 5, abnormal linkage was found between chromosomes 3 and 4 in both the male and female crosses involving the mutant, as shown in the heat maps of Fig 3. Such linkage is often the signature of a structural rearrangement between the two concerned chromosomes. Because we know that chromosome 3 is involved in post-meiotic selection, this inter-chromosomal linkage may likely induce post-meiotic selection on chromosome 4 as well. This would imply that CO counting on chromosome 4 does not faithfully reflect what occurs during meiosis. As a result, we will omit chromosome 4 in the remaining analyses.

### CO numbers reveal interchromosomal effect (IC)

Above, we described the impact of the structural rearrangement on COs in chromosome 3 (intrachromosomal effect). Now, we tested whether the recombination suppression in chromosome 3 affected the recombination in the other chromosomes (interchromosomal effect). Given the linkage between chromosomes 3 and 4, we examine COs in chromosomes 1, 2, and 5. Indeed, since these showed no abnormal linkage (Fig 3), we can reasonably conclude that chromosomes 1, 2 and 5 are not affected by post-meiotic selection and, therefore, we can use their genetic data as an estimation of CO number. When comparing mutant and control, pairwise comparisons of genetic length reveal excess of COs in the mutant for chromosomes 1, 2, and 5 in male meiosis but not in female meiosis (Table 2). Such increased number of COs in chromosomes 1, 2, 5 associated with recombination suppression in chromosome 3 is a signature of the IC effect. The difference in behavior between male and female meioses may be related to the fact that numbers of COs in female are close to the minimum value imposed by the obligatory CO rule, reducing the IC effect. This hypothesis is consistent with the fact that genetic lengths in Fmut tend to be higher than in Fctr when chromosomes 1, 2, and 5 are pooled, although this effect is not statistically significant. COs increased significantly in Mmut compared to Mctr (p<10^−7^ when pooling all three chromosomes). This analysis leads us to argue that chromosomes 1, 2 and 5 are clearly compensating the recombination suppression occurring in chromosome 3 by additional COs. To determine whether the IC effect is targeted to particular regions of the chromosomes 1, 2, and 5, we performed statistical tests for pairwise comparisons of the recombination landscapes of these chromosomes between control and mutant. This analysis did not reveal any significant difference except in the pericentromeric regions of chromosomes 2 (female meiosis) and 5 (male meiosis) (S2 Fig).

**TABLE 2.** CO numbers on chromosomes 1, 2 and 5 in the recombinant populations.

|  | Map size (cM) |  | Recombination Rate (cM/Mbp) |  | Proba | Map size (cM) |  | Recombination Rate (cM/Mbp) |  | Proba |
| --- | --- | --- | --- | --- | --- | --- | --- | --- | --- | --- |
|  | Mctr | Mmut | Mctr | Mmut |  | Fctr | Fmut | Fctr | Fmut |  |
| **Chr 1** | 123.13 | 141.37 | 4.05 | 4.65 | 0.02 | 79.64 | 83.84 | 2.62 | 2.76 | 0.50 |
| **Chr 2** | 74.61 | 94.93 | 3.79 | 4.82 | 0.001 | 62.80 | 60.32 | 3.19 | 3.06 | 0.65 |
| **Chr 5** | 110.99 | 141.47 | 4.12 | 5.25 | 8.08E-05 | 68.86 | 75.12 | 2.55 | 2.79 | 0.29 |
| **ALL** | 102.91 | 125.92 | 3.98 | 4.90 | 8.19E-07 | 70.43 | 73.09 | 2.79 | 2.87 | 0.62 |

Proba: *p*-values associated to the H0 hypothesis that control and mutant populations have the same numbers of COs. ALL: pooled analysis of chromosomes 1, 2, and 5.

To understand whether the additional COs could be associated to a change in the interference we measured the interference strength by fitting the segregation data to the so-called Two-Pathway Gamma model using the CODA software [29]. As for the recombination rates, we restricted the analysis to chromosomes 1, 2, and 5. As can be seen from S1 Table, there is a trend whereby the mutants generally have lower interference strength than the controls. However, the low number of pairwise comparisons (six, coming from the two sexes for each of the three chromosomes) prevented us from rejecting the null hypothesis of no trend (one-sided p-value of 0.17 when using the pooled data).

## Discussion

In this study, we have shown in *Arabidopsis* that the failure of a single chromosome pair to recombine in meiosis, following changes in its structure, has far reaching effects on the other chromosome pairs. In our case, the structural rearrangement is a large pericentric inversion involving chromosome 3. The inversion is associated to *Atmcc1* mutation previously characterized as homozygous line [23]. The genetic maps obtained in this work by 143 SNP markers in populations of over 400 individuals provide evidence of strong recombination suppression in chromosome 3 in both male and female meiosis. As expected, post-meiotic selection (PMS) operates against unbalanced gametes carrying deficiency/duplication of genomic regions that are usually produced by a single CO (or an odd number of COs) localized within the inverted region [30]. PMS is very high in our populations, presumably as a consequence of the existence in *Arabidopsis* of a relationship between the size of the inversion and the proportion of unbalanced gametes as reported for other organisms [30]. For this reason, genetic data cannot provide an estimation of CO number for chromosome 3 in *Atmcc1*. However, the chiasma count in heterozygous *Atmcc1* allowed us to establish that a real CO suppression occurs in chromosome 3. In particular, we measured 1.11 chiasmata per cell in chromosome 3 corresponding to a decrease of 34% compared to the control. The obligate CO was maintained thereby ensuring the correct segregation. In literature, the chiasma number for chromosome 3 is reported to range from 1.72 to 2.04 in different *Arabidopsis* ecotypes [31] including C24 and Landsberg, which were used in this work. Accordingly, 2.14 COs per meiosis, on average, were found by Giraut and colleagues [27] in a Col x Landsberg cross for chromosome 3, using genetic data. Based on comparable chiasma number of chromosome 3 between homozygous *Atmcc1* (1.68) and wild type (1.54), previously published by our group [23], we can exclude that factors other than the structural rearrangement are causative of the recombination decrease observed in heterozygous *Atmcc1* (this work). The reduction/suppression of COs within inversions is well documented across several organisms [12, 13]. Different mechanisms are invoked to explain this CO suppression such as non-homologous synapsis (hetero-synapsis) [32] and asynapsis [16] between the normal and inverted regions. However, in this work recombination events such as double COs are detected in chromosome 3 suggesting that homo-synapsis occurs in the rearranged region. Moreover, homo-synapsis is indirectly confirmed by PMS associated with having an odd number of COs. As reported in other species [33], homo-synapsis can imply the formation of an inversion loop or, alternatively, the pairing of the region with inversion in the anti-parallel orientation associated with the pairing failure of the outside regions. The latter seems to be confirmed by our data looking at COs in individual plants each representing a meiosis. Indeed, COs located in the inverted region are never coexisting with COs in the outside regions. Furthermore, no plants with COs on both outside regions are found. This pairing behavior can account for the deficit of COs on chromosome 3. Looking at total number of COs, we confirmed that the loss of COs within chromosome 3 was not fully counterbalanced by an increase of COs in outside regions. In diverse *Arabidopsis* structural rearrangements (inversions and deletions), γ-rays induced, Ederveen and colleagues [18] evidenced a different recombination behavior since loss of COs in the rearranged regions was generally compensated for by increase in COs on the unaltered part of the same chromosome. Similar behavior was found in *C. elegans* individuals carrying an inversion [14]. The differences between our case and those described in literature could be related to the size of the structural rearrangement. Indeed, in our material, the inversion covers 80% of the length of chromosome 3, implying that the outside regions are very restricted in size. The size of these regions, lying between the inversion boundaries and the telomeres, may constrain the opportunities for COs events. Our observation that COs take place preferentially in the outside region with superior size (at South arm) is consistent with this hypothesis. Besides the size, chromatin structure of these regions can have also an influence on CO formation. For example, in *Drosophila*, recombination suppression was estimated to extend 2.5-3 Mbp beyond the heterozygous inversion boundaries [20] and similar observations were reported in other organisms [34, 35]. Regarding telomeres, in budding yeast the 20 kbp region adjacent to telomeres exhibits a significant lower recombination rate [36]. In *Arabidopsis*, the telomeres are estimated to be 2-5 Kbp in length [37] but recombination data in these chromosomal domains are not available due to low resolution means of detecting recombination events in plants.

Perhaps the most striking aspect of this study is the finding that the failure of chromosome 3 to recombine is coupled to increased CO frequencies on the other chromosome pairs. Furthermore, we saw that this increase is associated with lower interference, although our limited data did not lead to a statistically significant difference. *A posteriori*, such a lower interference can be justified as the mechanism regulating CO number. Indeed, when going from DSB to COs, interference is commonly considered to be a way to ensure the presence of at least one CO while at the same time limiting the total number of COs. Reducing CO interference is then expected to lead to additional COs. Clearly, the enhancement in CO number on the other chromosomes (1, 2 and 5) can be thought of a compensation for the CO suppression existing on chromosome 3. This expectation is supported by our observation that CO number per cell is not affected in heterozygous *Atmcc1*. These compensatory increases are strictly an interchromosomal (IC) effect as was first described in *Drosophila* female meiosis [38, 39]. Recent examples of IC in flies carrying heterozygous inversion may be found in the works of Stevison and colleagues [20] and Joyce and McKim [40]. In the latter work, the authors suggested that heterozygous inversions cause a discontinuity in the alignment between homologs which can result in defects of axis structure. Consequently, the chromosome pair is destabilized over time, thereby inducing a pachytene delay that increases the chance of DSBs to become COs at the expense of NCOs. In this model, the global response of all the chromosome pairs to maintain the expected number of COs/cell is mediated by AAA+ ATPase PCH2 apparently without increase of DSB number. In *C. elegans*, asynapsis of the X chromosome resulted in IC response on autosomes [41]. In this case, IC associated to delay in meiotic progression, similarly to what happens in flies, requires, however, additional DSBs and their resolution as COs. In both flies and worm, IC is explained as an effect of the disruption of the normal timing of meiotic prophase events. To ensure the expected CO number/cell, based on our results we can suggest that also *Arabidopsis* has a window of opportunity for CO formation through either initiation of new DSBs or NCOs being redirected into COs. How a change on just one chromosome triggers the global response and the relative signaling pathway in *Arabidopsis* remains to be determined.

## Materials and Methods

### Plant material

The *Arabidopsis* genotypes used in this work include *Atmcc1* mutant, and the ecotypes C24 and *Landsberg erecta* (Ler). *Atmcc1* was previously isolated from an enhancer activation tagging population [22] obtained after floral transformation by *Agrobacterium tumefaciens* with the binary vector pSKI015 [42]. Homozygous line of *Atmcc1* is already characterized [23]. C24 is the genetic background of *Atmcc1*. C24 genotype used in this work has a T-DNA insertion on chromosome 3 between ABI3 and GL1 markers (S3 Fig) [43]. *Atmcc1* has an additional T-DNA insertion on the long arm of chromosome 3, at a location close to north telomere (S3 Fig). The accession of Ler (213AV) was kindly given by the Centre de Resources Biologiques at the Institut Jean Pierre Bourgin (Versailles, France) (http://dbsgap.versailles.inra.fr/vnat/). Plants were grown in controlled growth chambers with 16 h/8 h of light/dark at 22°C/18°C.

### Recombinant population construction

Ler was crossed with C24 and *Atmcc1* to obtain two F1 progenies. About 50 F1 hybrids were backcrossed with Ler using F1 plants either as the male or as the female parent to obtain four BC1 populations (Fig 1). For each BC1 population, about 700 seeds were sown *in vitro*. Subsequently, after two weeks, 430 seedlings transferred in pots were grown in the controlled growth chambers for three weeks.

### Genomic DNA extraction

DNA extraction was performed as described by Giraut and colleagues [27] with some modifications. Two hundred mg of green tissue from each BC1 plant was collected, freeze-dried, lyophilized and ground in 96 well plates closing hermetically the wells with plastic caps. One ml of Extraction Buffer (Tris pH 8 0.1 M, EDTA 50 mM, NaCl 0.5 M, SDS 1.25%, PVP 40000 1%, Sodium Bisulfite 1%, pre-warmed at 65°C) was then added to each well and the plates were incubated at 65°C for 30 min. Three hundred ml of cold 60% KAc 3 M, 11.5% glacial acetic acid was added to each well. The plate was sealed with a Thermowell film (Corning), shaken gently and placed on ice for 5 min. After centrifugation in a A-4-62 rotor (Eppendorf) at 3.220 g for 10 min at 4°C, 800 ml of the supernatant was transferred to a clean DeepWell plate and 1 mL of CGE buffer (1/ 3 Guanidine hydrochloride 7.8 M, 2/3 ethanol 96%) was added per well. 600 mL of the mixture was filtered with a Whatman Unifilter 800 GF/B plate placed on Deep Well plate (Greiner Bio-One) and centrifuged for 2 min at 5.806 g in a Nr 09100F rotor (Sigma) at room temperature. The flow-through was discarded. This step was repeated twice. The membrane was washed twice by adding 500 ml of Washing buffer (37% Aqueous solution, 63% ethanol 96%) (Aqueous solution: KAc 160 mM, Tris HCl pH 8 22.5 mM, EDTA 0.1 mM) and then centrifuged for 2 min at 5.806 g at room temperature. The DNA was eluted with 70 ml of H_2_O by centrifugation for 2 min at 363 g at room temperature. This step was repeated twice. RNAse A was added to 0.5 mg/ml and the DNA concentration was determined using the Quant-iT dsDNA BR assay Kit (Invitrogen) with an ABI 7900HT real-time PCR system (Applied Biosystems, MA, USA).

### Genotyping Single Nucleotide Polymorphism (SNP) markers

A genome-wide set of 146 SNP markers that are polymorphic between C24 and Ler was selected using POLYMORPH website (http://polymorph.weigelworld.org/cgi-bin/webapp.cgi?page=Home;project=MPICao2010) (Zeller et al. 2008, Clark et al. 2007). Twenty percent of SNPs were validated by SANGER sequencing in our lab. Physical position of each marker was verified using SEQVIEWER tool from www.arabidopsis.org. Evenly distributed informative SNPs with an average spacing of 0.8 Mb were used for genotyping BC1 populations. For each marker, a total number of 1680 plants was genotyped by LGC Genomics through KASP™ technology (https://www.lgcgroup.com/). DNA was purified with Whatman Unifilter plates and quantified to a master dilution of 5μg total DNA. DNA was transferred into appropriate microtiter plates using a LGC’s RepliKator™ robot. Final DNA concentration was 50ng/μl per well of sample and KASP procedure was carried out by LGC genomics following standard KASP assay procedure [44]. Alleles were called by KBioscience (S4 Fig). Each SNP was checked using SNPviewer2 and rescored whenever any error was observed in the clustering of the homozygous and heterozygous genotypes. Three SNPs were removed from the final dataset. Plants with more than 50% of uncalled or ambiguous data were discarded. Due to the high quality of the experiment, the total of eliminated plants is 21. The resulting BC1 populations comprised 418 plants in Mmut, 417 in Fmut, 414 in Mctr and 410 in Fctr. The list of 143 SNP markers are listed in S2 table and raw data of genotyping are reported in S3 table.

### Analysis of segregation distortion

To test whether any deviation from normal segregation (1:1) occurs in BC1 populations, the number of plants with the C24 allele at each marker locus was counted considering that the frequency of each allele should not deviate from 50% plus a delta value calculated as standard deviation according to Giraut and colleagues [27]. No significant segregation distortions were found in the populations except for few slight divergences that do not bias our estimate of genetic lengths (S5 Fig).

### Construction of Genetic Maps, Marey Maps and Linkage Heat Maps

The order of the markers was taken from the Weigel database [25, 26] for Ler. Nevertheless, for sake of completeness, we also determined it *ab initio* based on genetic linkage only. In the case of chromosome 3 in crosses involving the mutant, we found that the *ab initio* orders were ambiguous, reflecting the presence of the inversion. In contrast, for all other chromosomes, the *ab initio* orders were unambiguous and identical to the ones in the reference genome. For each BC1 population, genetic lengths between adjacent markers were calculated using Kosambi’s function. Based on these genetic maps we computed the associated Marey maps. Finally, linkage heat maps were produced by calculating the LOD for linkage for each pair of markers, be they on the same chromosome or not. Two-point linkage analyses were performed using the CarthaGene software [45] called from R scripts. Such heat maps can provide signatures of structural rearrangements within chromosomes and even reveal unexpected linkage between different chromosomes.

### COs and distribution of CO numbers per chromosome

For each BC plant, the number of COs was calculated by scoring allele changes across single chromosomes. Average number of COs per bivalent was estimated by dividing the total number of COs for that chromosome by the number of plants in the population and multiplying by 2.

### Cytological analysis of chiasmata

Male meiosis was investigated using the spreading technique described by Ross et al. [46] with some modifications. Briefly, floral buds, previously fixed in ethanol/acetic acid (3:1 v/v), were rinsed in freshly made ethanol/acetic acid (3:1 v/v), followed by citrate buffer. Then buds were digested in an enzyme mixture consisting of 0.3% (w/v) cellulase and 0.3% (w/v) pectolyase for 1h and 15’ at room temperature, followed by 30’ at 37°C. After digestion, the buds were rinsed in citrate buffer. One or two whole buds were then transferred on a slide and were macerated with a needle in 10 μL of 60% acetic acid on a hot plate (45°C) for 30s to release meiocytes. The meiocytes were then fixed with ice-cold ethanol/acetic acid (3:1 v/v). The slides were air dried and stained using 4’-6-diamidino-2 phenylindole (DAPI). Slides were analyzed by fluorescence microscopy. Chiasmata were recorded at metaphase I in PMCs after fluorescence in situ hybridization (FISH) following the method of Sanchez-Moran and colleagues [28].

## Supporting information

**S1 Fig. Relationship between physical and genetic positions, and corresponding recombination rate per chromosome.** X-axis: physical position (Mbp) of the SNPs. Left Y-axis and blue lines: genetic position (cM) of SNPs on Fctr (Female Control) and Fmut (Female Mutant) linkage maps. Right Y-axis and red lines: recombination rate (cM/Mbp) given by the derivative of the smoothed Marey map curve.

**S2 Fig. Illustration of the statistical test used to compare recombination landscapes between control and mutant populations for chromosomes 1, 2, and 5.** Solid curves: Marey maps normalized to the average genetic length, so the comparison focuses on differences in the shape of the recombination landscapes and is not affected by differences in the values of chromosome genetic lengths. Dotted curves: first derivative of the normalized Marey maps, indicating the recombination landscape along the chromosome. The black rectangles show the ten bins used for the analysis (from left to right: bin 1 to 10). Bin boundaries were chosen so each bin contained regions of the same genetic length. On top of each bar, a black vertical arrow indicates the difference between both populations in average recombination rates over the bin considered, and the error bars indicate the 95% confidence intervals of these average recombination rates. *p*-values corresponding to the H0 hypothesis that both populations have the same recombination landscape are indicated below the X-axis label.

**S3 Fig. Schematic representation of chromosome 3 indicating some random loci on short and long arm, centromere (blu) and T-DNA insertions (orange) in C24 and in *Atmcc1***. The first insertion (T-DNA 1) is present in both C24 and *Atmcc1* while the second insertion (T-DNA 2) occurs only in *Atmcc1*.

**S4 Fig. Example of KASP genotyping output.** Red dots represent Ler homozygous plants, green dots represent Ler/C24 heterozygous plants. Each plate contains blank control (black dot) and may contain ambiguous genotype score (pink dots).

**S5 Fig. Frequency of C24 allele (in orange) at each marker per chromosome in the mapping of Mctrl population.** Lines (grey) represent 1% confidence intervals of the expected 0.5 value under Mendelian segregation.

**S1 Table: Interference strength measured by fitting the data to a two-pathways Gamma model.** Nu: interference intensity in the interfering pathway (Class I crossovers). p: proportion of crossovers formed via the non-interfering pathway (Class II crossovers). nu_Inf, nu_Sup, p_Inf, p_Sup: lower (Inf) and upper (Sup) boundaries of 95% confidence intervals for nu and p, based on 1000 resimulations (see Materials and Methods).

**S2 Table: List of SNPs used for genotyping**.

**S3 Table: Genotyping scores of the recombinant BC1 populations**. 0 value represents Ler marker at homozygous state while 1 value is the presence of C24 marker.

## Acknowledgments

The authors thank Dr. C. Mezard and Prof. A. Barone for valuable suggestions and support about SNP genotyping. The authors are grateful to R. Nocerino for technical assistance in plant growth condition control.

## Author Contributions

### Conceptualization

Clara Conicella, Federica Consiglio, Matthieu Falque, Olivier Martin

### Formal analysis

Pasquale Termolino, Matthieu Falque

### Funding acquisition

Clara Conicella

### Investigation

Pasquale Termolino, Matthieu Falque

### Methodology

Gaetana Cremona, Antoine Ederveen, Rosa Paparo, Pasquale Termolino

### Project administration

Clara Conicella

### Software

Pasquale Termolino, Matthieu Falque

### Supervision

Clara Conicella, Federica Consiglio

### Validation

Pasquale Termolino

### Writing – original draft

Clara Conicella, Pasquale Termolino

### Writing – review & editing

Clara Conicella, Federica Consiglio, Matthieu Falque, Olivier Martin, Pasquale Termolino

